# Modulation of Cysteine modifications upon short time T-cell activation

**DOI:** 10.64898/2026.01.11.698897

**Authors:** Komal K Mandal, Peter Jensen, Martin R Larsen

## Abstract

T cell activation is initiated by rapid and tightly regulated signaling events that rely on reversible post-translational modifications to ensure speed, specificity, and fidelity. While phosphorylation is well established as a central regulator of T cell receptor signaling, the contribution of cysteine-based modifications remains poorly defined at a systems level. Due to their redox-sensitive thiol side chains, cysteine residues can undergo rapid and reversible chemical modifications, enabling them to function as molecular switches that modulate protein structure, activity, and interactions during signal transduction. Engagement of the TCR induces localized redox signaling, primarily through mitochondrial and NADPH oxidase-derived H₂O₂, leading to selective and reversible oxidation of reactive cysteine residues. These redox events act as spatially and temporally controlled signaling mechanisms rather than global oxidative stress, transiently reshaping phosphorylation-dependent networks through modulation of redox-sensitive kinases, phosphatases, and adaptor proteins. Specificity is conferred by peroxiredoxin-, thioredoxin-, and glutathione-dependent redox control systems that fine-tune cysteine reactivity. In addition, redox-dependent modulation of cysteine residues within zinc-coordinating motifs suggests a coupled redox–zinc signaling axis during T cell activation, where transient redistribution of zinc from cysteine-rich zinc-finger proteins may further regulate kinase activity and transcriptional programs downstream of TCR engagement.

Here, we present a quantitative, time-resolved proteomic strategy to systematically interrogate cysteine dynamics during the earliest phases of human T cell activation. By combining cysteine-specific phosphonate adaptable tagging, thiol–disulfide exchange chromatography and TMT-based multiplexed mass spectrometry, we quantified 25,324 free cysteine-containing peptides and 14,853 reversibly modified cysteine peptides across seconds-to-minutes following TCR stimulation. This comprehensive cysteine proteome revealed rapid remodeling of cysteine redox states as early as 30 seconds after activation of T-cells. Functionally, regulated cysteines were enriched in core signaling pathways, including kinases, phosphatases, transcription factors, and nuclear zinc-finger proteins. Notably, key TCR signaling components such as Lck, CD45, ZAP-70, Akt, NFATC3, and RASGRP1 exhibited dynamic cysteine modulation. Together, these findings establish cysteine redox dynamics as a pervasive and functionally relevant regulatory layer in T cell activation, positioning cysteine modification alongside phosphorylation as a central component of immune signal transduction.

## Introduction

Cysteine (Cys) is a highly conserved amino acid characterized by a chemically labile and redox-sensitive thiol side chain. Conserved cysteine residues are frequently found in enzyme active sites or within solvent-exposed regions of proteins, where they are well positioned to participate in catalysis, metal coordination, and protein–protein interactions (1, 2). The reactive sulfur atom of the cysteine side chain enables the formation of irreversible covalent modifications, such as sulfonic acid, as well as a broad range of reversible oxidative modifications, including S-nitrosylation, S-sulfenylation, S-glutathionylation, S-sulfhydration, S-acylation, and disulfide bond formation (1). Through these reversible and irreversible chemistries, cysteine residues acquire diverse functional roles, most prominently in disulfide bond formation and metal binding (3).

One of the most important metal-mediated functions of cysteine is its ability to coordinate zinc ions. Arrays of cysteine residues form structural motifs such as zinc fingers and zinc clasps, in which Zn²⁺ is coordinated by cysteine and/or histidine side chains to stabilize protein folds and enable sequence-specific interactions with DNA, RNA, or other proteins (4, 5). These zinc-binding motifs are particularly abundant in transcription factors, chromatin regulators, and signaling adaptors, positioning cysteine-zinc conjugation as a critical mechanism for controlling protein structure and function. Importantly, zinc coordination by cysteine is inherently redox-sensitive; oxidative modification of coordinating cysteines can weaken zinc binding, alter protein conformation, and modulate downstream biological activity (6, 7).

Cysteine modifications can occur rapidly and reversibly in response to subtle changes in cellular conditions, including fluctuations in pH or redox state. These properties enable cysteine-containing proteins to function as oxidases, reductases, peroxidases, disulfide isomerases, and transcription factors, making cysteine residues central regulators of cellular redox homeostasis (8). The ability of cysteine to act as both a sensor and a molecular switch, detecting environmental cues and reversibly transitioning between functional states, highlights its potential importance in signal transduction processes (8).

In immune cells, redox signaling - particularly superoxide-mediated signaling - is well characterized (9). However, redox-based regulation extends beyond superoxide neutralization and can influence signaling through multiple molecular mechanisms. Upon short-time stimulation, such as antigen engagement of immune receptors, localized production of reactive oxygen species (ROS), especially hydrogen peroxide (H₂O₂), occurs (10–12). Due to its relative stability and diffusibility, H₂O₂ can function as a second messenger that selectively and reversibly oxidizes reactive cysteine residues. These localized redox events generate spatially and temporally restricted redox microenvironments that enable rapid modulation of cysteine reactivity within seconds of stimulation. In addition to direct oxidation, cysteine states can be dynamically reshaped through glutathione-dependent reactions, thioredoxin- and peroxiredoxin-mediated redox relays, and redox-dependent changes in zinc coordination, collectively providing multiple layers of regulation following receptor activation.

In T cell signaling, the non-kinase T cell receptor (TCR) initiates activation upon antigen binding but relies on the cytoplasmic kinase Lck to transmit activation signals into the cell (13, 14). Lck phosphorylates the cytoplasmic tails of the TCR complex and activates the downstream kinase ZAP-70, which then binds to the phosphorylated TCR and propagates the activation signal (13, 14). Notably, cysteine residues within the Src homology domains of ZAP-70 have been shown to regulate phosphotyrosine binding and signal propagation, highlighting a direct role for cysteine chemistry in TCR signaling (15). Moreover, immune receptors, including immunoglobulins and integrins, are organized as multimeric complexes stabilized by disulfide bonds and zinc-coordinated structures that function as redox-sensitive switches to regulate receptor activity.

Together, these observations suggest that cysteine chemistry, through reversible oxidation, disulfide exchange, and zinc coordination, constitutes an important but incompletely understood regulatory layer in T cell signaling. Comprehensive, large-scale characterization of cysteine-mediated regulation offers a powerful alternative to conventional fluorescence-based or genetic perturbation approaches and has the potential to reveal previously unrecognized mechanisms of immune signal transduction (16). Recent advances in mass spectrometry-based proteomics have enabled high-throughput interrogation of signaling networks. However, due to the frequent low stoichiometry of cysteine modifications, selective enrichment of cysteine-modified peptides is required. One such approach involves tagging cysteine residues with phosphonate-based adaptable tags (PATs), enabling selective enrichment using TiO₂ chromatography (17). The high affinity of TiO₂ for phosphate-containing moieties is well established (18, 19). In addition, thiol–disulfide exchange (TDE) chromatography has been employed to selectively enrich peptides containing reversibly modified cysteines (20) and sodium deoxycholoate acid precipitation (SDC-ACE) has recently been used for enrichment of S-palmitoylated peptides (21). The methods for enriching cysteine containing peptides are frequently combined with various selective reduction reagents to selectively target one or more cysteine modifications (22).

Here we combined the CysPAT strategy in combination with TiO_2_ enrichment (17, 18), with the TDE chromatography to explore the modulation of free Cysteines (freeCys), as well as reversibly modified Cysteines (RmCys) in the very early signalling events of the T cell activation. We demonstrate that cysteine residues undergo extensive and rapid modulation during the earliest phases of TCR activation. By quantitatively profiling both freeCys and RmCys, we reveal that cysteine chemistry represents multiple, largely distinct regulatory layers within the signalling network. Core components of the TCR signalling cascade, including Lck, CD45, ZAP-70, and Akt, exhibit dynamic cysteine regulation within seconds/minutes of stimulation. In addition to signalling kinases and phosphatases, cysteine modulation targets transcription factors and RNA-processing proteins, indicating early coordination between signalling and gene regulatory programs. A prominent enrichment of zinc-coordinating cysteines and zinc-finger proteins suggests a coupled redox–zinc regulatory axis operating during T cell activation. These findings support a model in which localized redox signalling rapidly reshapes cysteine reactivity through oxidation, disulfide exchange, and zinc coordination. Together, our work establishes cysteine-based regulation as a pervasive and functionally relevant mechanism in early T cell signalling.

## Materials and methods

All chemicals were obtained from Sigma Aldrich, unless otherwise is stated.

### Cell culture

Jurkat clone E6-1 was grown in RPMI enriched with 10% fetal bovine serum (FBS) (v/v), 10mM HEPES, penicillin (100units/ml) / streptomycin (100µg/ml), 5%CO_2_ at 37°C, till it reached confluency (no more than 1.5×10^6^cells/ml). The cells were counted using trypan blue, this stain was also used to verify the viability of cells.

### T cell activation by antiCD3 and antiCD28 antibodies

The cells were serum starved for 9hrs. After starvation, the cells were washed once in RPMI without serum and subsequently resuspended in RPMI with 10mM HEPES. Approximately 5×10^7^ cells were used for each activation time point in a volume of 100 µL. Cells were activated by adding, 1:1 pre-crosslinked mixture of anti-CD3 (Santa Cruz, clone HIT3a) and anti-CD28 (Santa Cruz, clone CD28.2) antibody. The crosslinking antibody was Goat anti-Mouse IgG (Thermos Fisher). Activation time points were: 0s, 30s, 60s and 300s, and each activation were performed in quadruplicate at 37°C. After completion of the activation at the precise timepoints, the samples were snap frozen in liquid nitrogen. All the samples were stored in -80°C until further processing.

### Protein extraction

To each snap-frozen sample SDS was added to 1% and Neocuproine was added to a final of 0.1 mM and the samples were boiled for 5 min at 95°C. After boiling, each sample was sonicated, which involved three on/off cycle for 20sec each with 60% amplitude settings via probe tip Q125 Sonicator (Qsonia), to ensure total cell lysis and DNA degradation. After sonication, the samples were centrifuged at 20.000g for 20 min and the supernatant was transferred to another low binding Eppendorf tube. Protein concentration was measured using an Implen NanoPhotometer N60 (Implen, Germany).

### Western Blotting

To confirm the T cell activation, aliquot of 25µg proteins from each time point was blotted for Zap70, Zap70 [p-Y493] (#2704 Cell Signaling Technology), CD247/CD3ζ [p-Y72] (MA5-28537 ThermoFisher Scientific); CD247/CD3ζ [p-Y142] (PA5-37512 ThermoFisher Scientific) and ß-actin (Abcam). Briefly, SDS PAGE separation of proteins was performed using 4-12% Bis-Tris plus gels (Invitrogen), followed by transfer of the proteins to a 0.45-micron PVDF immobilon transfer membrane (Merck) using a Transblot SD semi dry transfer cell blotter system (Bio-Rad). The membranes were blocked in 3% BSA in Tris Buffered saline (TBS) at room temperature with gentle shaking for 1hour. Incubated with primary antibodies was performed overnight at room temperature. After incubation, the membranes were washed with 50ml of TBS three times, each for 5 minutes. HRP-conjugated secondary antibody was prepared in TBS and the membrane was incubated with this antibody for 2 hours. The membrane was washed with 50ml of TBS three times. Signals were developed using chemiluminescence system (Luminata Forte Western HRP substrate, Millipore), and the bands were imaged using semi-automatic settings in an Amersham imager 680 (GE). The image was stored as 8bit images, and the blots were analyzed using the image J tool.

Note: concentration of antibodies according to manufacturer’s recommendation, secondary antibodies were Goat-anti-rabbit IgG, HRP-linked (# 7074, Cell Signaling), rabbit anti-mouse IgG H&L, HRP linked (ab6728, Abcam).

### CysPAT tagging of free cysteines and buffer exchange

The CysPAT tag for 16 samples was prepared in the following way: 4mg SIA (14.2 μmol) was dissolved in 20μl of dimethyl sulfoxide (DMSO) and 4mg (32 µmol) of 2-aminoethylphosphonic acid (2-AEP) was dissolved in 320μl of 100 mM TEAB buffer, pH 8 (Sigma T7408). The two reagents were mixed slowly by adding the aqueous 2-AEP solution drop by drop to the DMSO containing SIA solution during vortexing. After mixing, the sample was adjusted to pH 8 using 1M TEAB, and the final volume was made up to 565μl using 100mM TEAB buffer, pH 8, to ensure that the final concentration of the synthesized CysPAT is 25mM. The reaction is allowed to happen in the dark for 2 hours on a rotor at room temperature (RT). The tags are always prepared fresh and used immediately (17). A total of 200µg of protein from each of the 16 samples were incubated with CysPAT tags (final concentration of 5 mM) for 2 hours in the dark while rotating to covalently modify the free cysteines with a phosphonate group. After incubation, the samples were subjected to washing on a 10 kDa spin filter (Millipore) to remove excessive CysPAT reagent and to replace SDS with 1% Sodium Deoxycholate (SDC). After centrifugation, the samples were washed twice with 450 µL 2% SDC in 100 mM HEPES, pH 8.5. After the final wash, the solutions were transferred to low binding Eppendorf tubes and the samples were adjusted to 1% SDC, 100 mM HEPES, pH 8.5 containing 5 mM TCEP for reduction of reversibly modified Cysteines. The samples were incubated at room temperature for 20 min at RT.

### Trypsin/Lys-C digestion and TMT labelling

A total of 100μg of proteins was digested by adding 2µl of Endoprotease Lys C (0.01U) (Wako) to each sample and incubated for 1 hour at 37°C. After incubation, in-house generated methylated trypsin (23) was added to 4% (w/w) and subsequently incubated overnight at 37°C. After digestion, the peptides from each sample were labelled with TMTpro™ 16plex tandem mass tags (TMT) (Thermo Fisher) by carefully following manufacturer’s protocol. The labelling strategy were as follows: 0sec: 126, 127N, 127C, 128N; 30sec: 128C, 129N, 129C, 130N; 60sec: 130C, 131N, 131C, 132N; 300sec: 132C,133N, 133C, 134N. For each peptide solution, the TMT tag was solubilized in 20 µL 100% Acetonitrile and added to the 100µg peptide sample. After mixing the pH was measured and adjusted to 8-8.5 prior to incubation at RT for 1.5 hours. After incubation, a 1µL aliquot of each sample were mixed in the same tube containing 20 µL H_2_O, formic acid was added to precipitate the SDC, the sample was centrifuged at 20000g for 10 min and approximately 0.5 µg was used for high resolution LC-MSMS for evaluation of TMT incorporation efficiency and labelling ratios. After testing, the TMT samples were mixed to 1:1 and the SDC present in the sample was precipitated by adding 2% formic acid (v/v) followed by centrifugation at 20000g for 10 min. The supernatant was transferred to another tube and lyophilized.

### Enrichment of Phosphopeptides and CysPAT tagged cysteine-peptides (free Cys) by TiO_2_ bead purification

TiO_2_ beads can simultaneous enrich CysPAT tagged peptides, phosphorylated peptides as well as sialylated N-linked glycopeptides using TiO_2_ chromatography (19, 24, 25). The dried peptides were resuspended in 1ml of TiO_2_ loading buffer (1M glycolic acid, 80% ACN (v/v) and 5% TFA(v/v)). A total of 6 mg of TiO_2_ beads (GL Sciences Inc.) were added and the samples were incubated with shaking for 15min. After incubation, the tubes were centrifuged, the supernatant was transferred to a new tube containing 3mg of TiO_2_ beads and was incubated with shaking for 15min. The tubes were centrifuged and the supernatant was transferred to a low binding Eppendorf tube and lyophilized (TiO_2_ FT). The TiO_2_ beads from both the tubes (6.0+3.0mg) were pooled and washed once with wash first with wash buffer 1 (80% ACN (v/v), 1% TFA (v/v)) followed by wash buffer 2 (10% ACN (v/v), 0.1% TFA (v/v)). The second wash (Wash 2) was lyophilized and analysed separately. The washing steps include adding 100µl wash buffer, vortexing for 10s followed by centrifugation to pellet the beads and removing the supernatant. The CysPAT tagged peptides, phosphopeptides and sialylated N-linked glycopeptides, bound to the TiO_2_ beads were eluted with 1% ammonium hydroxide solution (v/v), pH 11. The eluate passed over a C8 (3 M Bioanalytical Technologies) stage tip to trap any residual TiO_2_ beads. The TiO_2_ was subsequently washed with 50% ACN/1% ammonium hydroxide, centrifuged and the supernatant was passed over the same C8 Stage tip to recover bound peptides into the eluted peptide fraction. The eluted peptides were vacuum dried to remove all the traces of ammonia.

The peptides eluted from the TiO_2_ beads were solubilized in 200 µL 20 mM HEPES, pH 7.2 and 15 µL agarose conjugated phospho-tyrosine monoclonal antibody (P-Tyr-100, Cell Signaling Technology (CST-9419S)) was added to the solution and incubated for 1 hour at RT with rotation. After the incubation, the beads were pelleted by soft centrifugation, and the supernatant was transferred to another low binding Eppendorf tube. The beads were washed once with 50 µL 20 mM HEPES, pH 7.2 and the supernatant was merged with the other supernatant. The beads were washed twice with 100 µL H_2_O and the phosphor-tyrosine peptides were eluted with 100 µL 0.3% TFA and subsequently purified by a Poros R3 microcolumn as previously described (REF).

The supernatant was subjected to deglycosylation of N-linked sialylated deglycopeptides using 1µl of PNGase F (New England Biolabs) and 0.5µl of Sialidase A (ProZyme) over night at 37°C. After deglycosylation, the peptides were reduced and alkylated using 5 mM DTT for 15 min and subsequently 10 mM Iodoacetamide for 15 min. After alkylation, the sample was acidified using 10 µL 100% TFA and 600 µL 100% ACN was added followed by the addition of the same TiO_2_ beads as used above. This step allows the phosphopeptides and CysPAT peptides to be enriched from the deglycopeptides. The flow through from the second TiO_2_ enrichment was lyophilized and desalted using a Poros R3 microcolumn (Deglycopeptide fraction) and the phosphopeptides and CysPAT peptides were eluted from the TiO_2_ as described above and lyophilized (Phospho/CysPAT fraction). The Wash 2 from the TiO_2_ was deglycosylated as above.

### Isolation of reversibly modified cysteine by thiol–disulfide exchange (TDE) chromatography

The peptides from the first TiO_2_ flow through (TiO_2_ FT) were resolubilized in 1 mL 0.1% TFA and desalted on a HLB column as described previously (26). After elution with 60% ACN in water and lyophilization, the desalted peptides were resolubilized in 100 µL 250 mM HEPES, pH 8, containing 3 mM TCEP for reduction of disulfide bridges, and incubated at 37°C for 30 min. After incubation, the sample was diluted to 1 mL in UHQ water and 150 µL S3 high-capacity acyl-rac capture beads (NANOCS Inc, cat. no. AR-S3-2) was added and incubated with rotation for 1 hour at RT. After incubation the beads were washed 4 times, once with 500 µL 1% SDS and 3 times with 500 µL H_2_O. All washing steps were performed in a MobiSpin column filter. To release the cysteine-containing peptides from the beads 300 µL of 100 mM HEPES, pH 8, containing 20 mM DTT was added to the beads on the filter and incubated at 37°C for 30 min. The filters were centrifuged and the solution containing the reversibly modified cysteine-containing peptides was collect in a low-binding Eppendorf tube, labelled “RmCys peptides”. The beads were washed using 100 µL of 100 mM HEPES, 20 mM DTT, pH 8, and collect the solution in the “RmCys peptides” Eppendorf tube. Finally, 45 mM iodoacetamide was added to the solution and it was incubated at RT in the dark for 30 min. After incubation, the peptide solution was desalted using a Poros Oligo R3 microcolumn as previously described (26) and eluted peptides were lyophilized.

### Pre-fractionation by high pH Reversed Phase (RP) chromatography

To reduce the complexity of the samples, offline high-pH RP liquid chromatography (HpH) was performed. A Dionex Ultimate 3000 system (Thermo Scientific) equipped with C18 analytical column (300µM × 100 mM), 1.5μM particle size (Waters) and executed by the Chromeleon framework, was used. The mobile phase comprised of: Solvent A (20mM ammonium formate, pH 9.3) and Solvent B (80% ACN (v/v) + 20% solvent A (v/v)). The Phospho/CysPAT fraction and RmCys peptide fraction were resuspended in 30 µl HpH solvent A and fractionated in a linear gradient from 2-50% B in 103 min, 50-95% B up to 108min, an additional 10min on constant 95% B followed by 15min in 2% B. The peptides were separated into a 96 well plate as 20 concatenated fractions. For the deglycopeptide fraction and Wash 2 fraction, the separation was performed into 10 concatenated fractions using a gradient from 2-50% B in 60 min. On completion, the plates were vacuum dried and stored in -20°C until further use.

### Reversed phase nano-LC_ESI-MS/MS

All peptides were reconstituted in 0.1% formic acid (FA). The samples were loaded onto a 25cm Aurora series emitter column (Ionopticks) with 75µm ID and packed with 1.6µm C18 via an Easy-nLC system 1000 (Thermo Fisher Scientific). During operation, the column was maintained at a constant temperature of 45°C in a homemade column oven. The Easy-nLC was connected to an Orbitrap Lumos (Thermo Fisher Scientific) equipped with a FAIMS ion mobility device. The A-buffer was 0.1% Formic acid, and the B-buffer was 95% ACN in 0.1% Formic acid. The tyrosine-phosphopeptide sample was separated using a 140min gradient beginning from 4-28% B for 110min, 28–50% B for 30min, 50–95% B for 2min, for 10min at 95% B to the wash the column followed by equilibration from 95-4% B within 1min and continue at 4% B for 6 minutes, at a constant flow of 400 nL/min. For all the other fractions, each fraction was separated using a 120 min gradient beginning from 4-28% B for 100min, 28–50% B for 20min, 50–95% B for 2min, for 10min at 95% B followed by equilibration from 95-4% B within 1min and continue at 4% B for 6 minutes, at a constant flow of 400 nL/min.

The spectrometer was set to positive ion mode using data-dependent acquisition and gained a full MS scan with an automatic gain control (AGC) target of 1x10^6^ ions and a maximum filling time of 50ms. Each MS scan was acquired at 120,000 full width half maximum (FWHM) resolution and a mass range of 350–1500 m/z. Field asymmetric ion mobility spectrometry (FAIMS) was used with two compensation voltages (CVs) of -50, and -70. The instrument was operated with a cycle time of 1.5 sec for each CV scan. Ions were selected for higher energy collision-induced dissociation (HCD) fragmentation with a collision energy of 40. Fragmentation was performed with 50,000 FWHM resolution for an AGC target of 3x10^4^ and a maximum injection time of 200ms using an isolation window of 1.2 m/z. Qual browser Thermo Xcalibur v4.1.31.9 was used to view the raw files.

### Data processing | Data normalization | Statistical analysis

The raw files were processed using Thermo Proteome Discoverer v2.5 (Thermo Scientific) with the database search program SEQUEST HT. The Uniprot Homo sapiens database (v2025, 26/02; 20574 entries) was used with following parameters: precursor mass tolerance 10ppm; fragment mass tolerance: 0.05Da; enzyme: trypsin; missed cleavage: 2; TMTpro was set as fixed modifications (N-term and K). Variable modifications for all the High pH fractions containing phosphopeptides, CysPAT and deglycopeptides were set to N-terminal acetylation (protein), SIA (C) (CysPAT modification), carbamidomethyl (C) (RmCys), Deamidation (N) (deglycopeptides) and phosphorylation (S/T/Y). For the RmCys fraction the variable modifications were set to SIA (C) and carbamidomethyl (C). The searched results were finally filtered to a threshold of 1% false discovery rate (FDR 0.01) using percolator software (27). The peptide-lists were exported to Excel. Statistical testing on the data was performed using Perseus platform the (v1.6.6.0) (28). For the time course, multiple ANOVA test with a 0.05 p-valve cutoff was applied and was further corrected using Benjamini-Hochberg correction factor for further 5% FDR (0.05). The peptides which passed this cutoff were selected for further processing and data mining. Qualitative analysis such as principal component analysis (PCA) were made for visualization. Euclidean based hierarchical clustering was made to view the expression trends along the time course.

### Analysis of the regulated peptides using bioinformatic tools

The ShinyGo application was used to perform Gene Ontology analysis (29). The STRING platform was used for visualizing protein networks (30). All the graphs were made using Microsoft Excel.

## Results and Discussion

### Experimental strategy with a focus on regulatory cysteine dynamics during early TCR activation

Dynamic post-translational modifications (PTMs) following T-cell activation have traditionally been studied primarily in the context of phosphorylation (14, 31) and ubiquitination (32). In contrast, the potential regulatory role of cysteine chemistry during the earliest stages of T-cell activation remains poorly understood (33). To address this, we investigated the temporal dynamics of free cysteines (freeCys; C–SH) and reversibly modified cysteines (RmCys; e.g., disulfides, S-nitrosylation, sulfenylation, mixed disulfides) during early TCR signaling.

To capture both proximal and more distal signaling events, we analyzed four activation time points: 0 s, 30 s, 60 s, and 300 s following antibody-mediated TCR stimulation. Signaling was terminated precisely at each time point by rapid snap-freezing in liquid nitrogen. Upon sample processing, the frozen samples were added SDS to a total of 1%, following 5 min boiling at 95°C and subsequent probe sonication. This procedure efficiently quenched enzymatic activity and preserved transient PTM states.

Our experimental workflow combined TMTpro-based multiplexed quantitative proteomics with a modified version of a previously established strategy enabling simultaneous analysis of phosphorylation, free cysteines, and reversibly modified cysteines (18). Free cysteines were labeled at the protein level using the CysPAT reagent immediately following T cell stimulation. After removal of excess CysPAT reagent, reversibly modified cysteines were reduced using TCEP. Proteins were digested and peptides labeled with TMTpro reagents, enabling multiplexed quantification of all conditions in quadruplicate. Phosphopeptides and CysPAT-labeled freeCys peptides were enriched using TiO₂ chromatography, while RmCys peptides were subsequently purified from the TiO₂ flow-through using TDE chromatography. The complete experimental strategy is illustrated in **Figure 1**.

**Figure 1:**
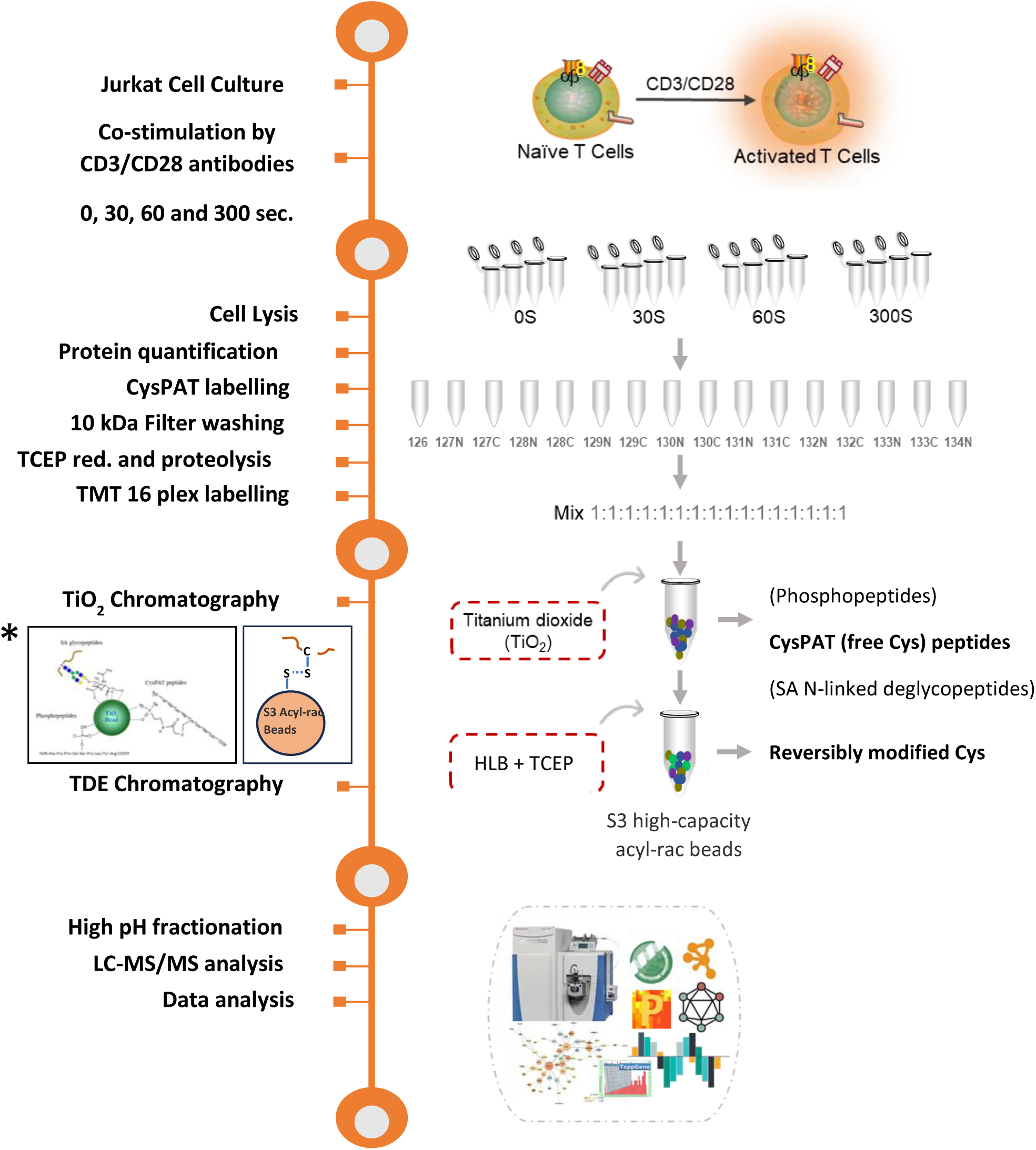
Illustration of experimental workflow with special focus on cysteine dynamics. Jurkat cells were activated for 0, 30, 60 and 300 seconds using a crosslinked antibody for CD3 and CD 28 and the activation was terminated by snap freezing in liquid Nitrogen. Cells were solubilized in 1% SDS, free cysteines on proteins were labelled with CysPAT tags. The excess tag was removed by 10 kDa filters and reversibly modified cysteines were reduced with TCEP. After protein digestion, peptides were labelled with TMT 16 plex and mixed. Phosphopeptides, sialylated (SA) N-linked glycopeptides and CysPAT peptides were enriched using TiO_2_ chromatography. The reduced RmCys peptides were purified using thiol beads (S3 acyl-rac beads). All peptide fractions were subjected to high-pH pre-fractionation. The peptides were subseqeuntly analysed using RP-nLC-MS/MS. After database searching the data was analyzed using several bioinformatic resources. *(insert) illustration of binding of TiO_2_ beads and S3 acyl-rac beads to respective peptide groups.

### Phosphoproteomics confirms robust and rapid TCR activation

Phosphoproteomic analysis was included to verify effective T-cell activation and to provide a reference framework for interpreting cysteine dynamics. The analysis revealed a total of 13038 unique phosphopeptide groups in 3610 phosphoproteins (**Figure S1A**). Of these phosphopeptide groups a total of 567, 1001 and 1783 were found differentially regulated upon TCR activation after 30 sec, 60 sec and 300 sec, respectively. To verify the activation of the TCR we investigated phosphorylation of the T-cell surface glycoprotein CD3 chains; CD3D, CD3E, CD3G and CD3Z. All CD3 chains contain immunoreceptor tyrosine-based activation motifs (ITAMs) in their cytoplasmic domain. Upon TCR activation, these ITAMs become phosphorylated by Src family protein tyrosine kinases LCK and FYN, resulting in the activation of downstream signaling pathways in the TCR signaling cascade (10, 14, 34–36). As evident from **Figure S1D**, we identified 2, 1, 2 and 6 unique phospho-tyrosine peptide groups in CD3D, CD3E, CD3G and CD3Z, respectively. Each of the phospho-tyrosine sites in the 4 proteins were localized to known ITAM domains. All phospho-tyrosine sites were significantly upregulated after all 3 timepoints (2-7-fold), indicating a proper T-cell activation after the crosslinked antibody activation.

### Overview of the identified free and reversibly modified Cys, their overlap and GO trends

The present strategy to investigate various Cys populations and their role of T cell activation resulted in quantification of 25,324 freeCys containing peptides (carrying CysPAT) and 14,853 peptides with RmCys (**Figure 2A**). These peptides could be annotated to 7100 and 5604 proteins, respectively. In total 5,813 and 1,560 freeCys and RmCys peptides were found to be differentially regulated at one of the three activation time points relative to the control based on ANOVA test and subsequent Benjamini-Hochberg correction, p<0.05 (**Figure 2A)**. Principal component analysis revealed clear temporal separation of cysteine states, with a pronounced divergence between unstimulated cells and those activated for only 30 seconds (**Supplementary Figure 2B and 2C**), highlighting the rapid and extensive remodelling of cysteine chemistry immediately following TCR engagement. Despite a modest overlap between regulated freeCys and RmCys peptides (8.1%), nearly 24% of regulated proteins were shared between the two fractions (**Figure 2B**), indicating that the same proteins often undergo multiple modes of cysteine regulation. The higher number of regulated freeCys peptides likely reflects stoichiometric considerations: free cysteines are generally less abundant than reversibly modified cysteines (many of which reside in stable disulfide bonds), making relative changes in freeCys more readily detectable.

**Figure 2:**
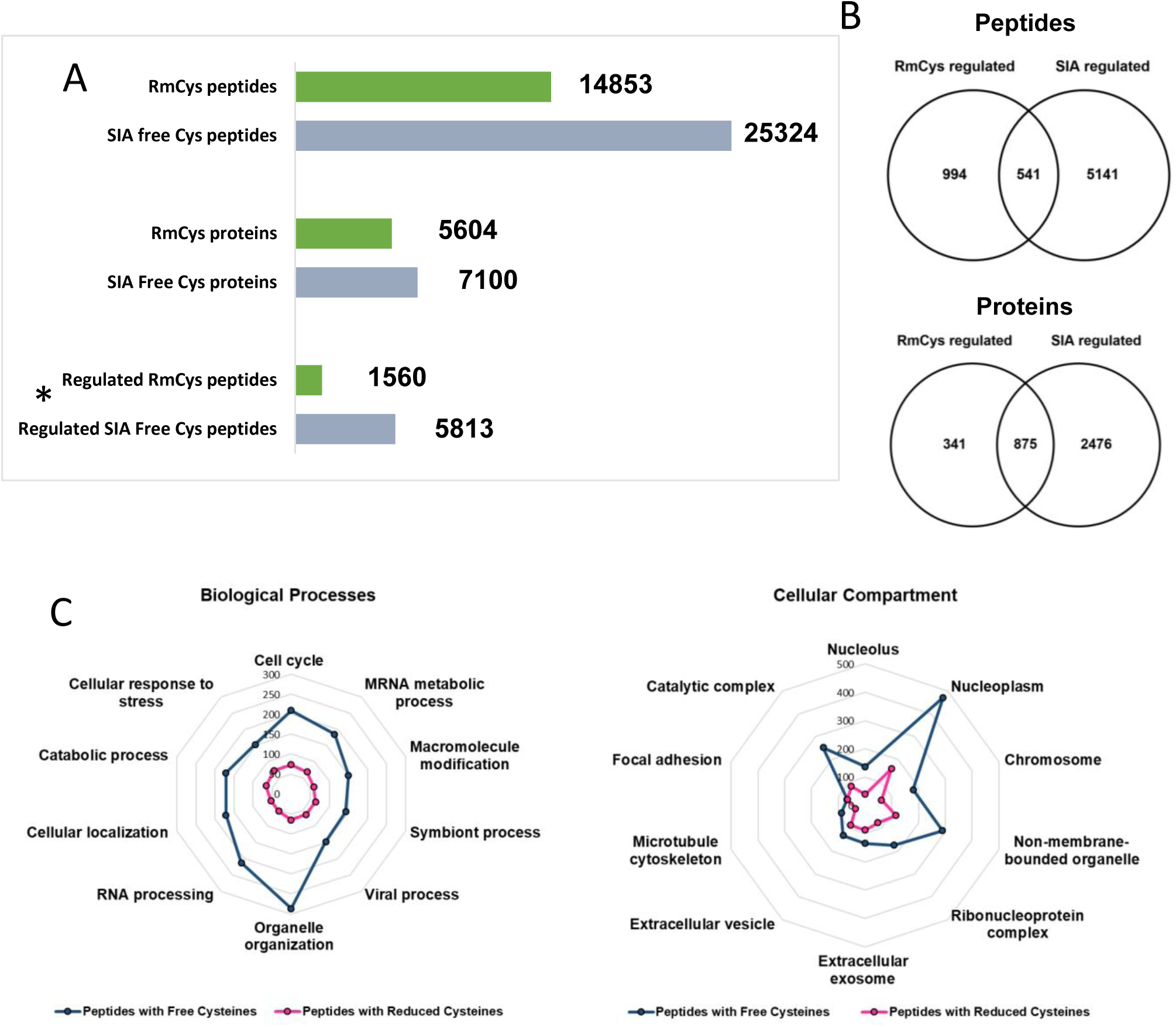
Descriptive overview of FreeCys (SIA) and RmCys proteins and peptides (A) Bar plots depicting the number of proteins and peptides identified with both kind of Cys including the number of regulated Cys in both categories based on ANOVA, Benjamini Hochberg p<0.05. (B) Venn diagram showing the overlap between regulated proteins and peptide groups with Free Cys and RmCys, respectively (C) ShinyGO app was used to evaluate the biological processes and cellular compartments enriched in each fraction, which revealed both the types of Cys were similar in their functions as well as localization in the cell.

Gene ontology analysis revealed highly similar biological themes for freeCys and RmCys proteins, including enrichment in catabolic processes, RNA processing, organelle organization, and nuclear localization (**Figure 2C**). Notably, freeCys proteins were disproportionately enriched in catalytic complexes, consistent with the known role of free cysteines in regulating enzymatic activity, particularly in redox-sensitive signaling enzymes such as phosphatases and ubiquitin ligases. The GO analysis is visualized as spider plots in **Figure 2C**.

### Early TCR activation induces rapid and structured cysteine remodeling across functional protein networks

To visualize the temporal dynamics of cysteine regulation following TCR activation, all significantly regulated peptides from both the freeCys and RmCys fractions were combined and subjected to hierarchical clustering analysis (**Figure 3**). This analysis resolved the peptides into five distinct clusters across the four activation time points. Notably, extensive cysteine modulation was already evident after only 30 seconds of stimulation, highlighting the rapid nature of cysteine-based regulation during early T-cell activation.

**Figure 3:**
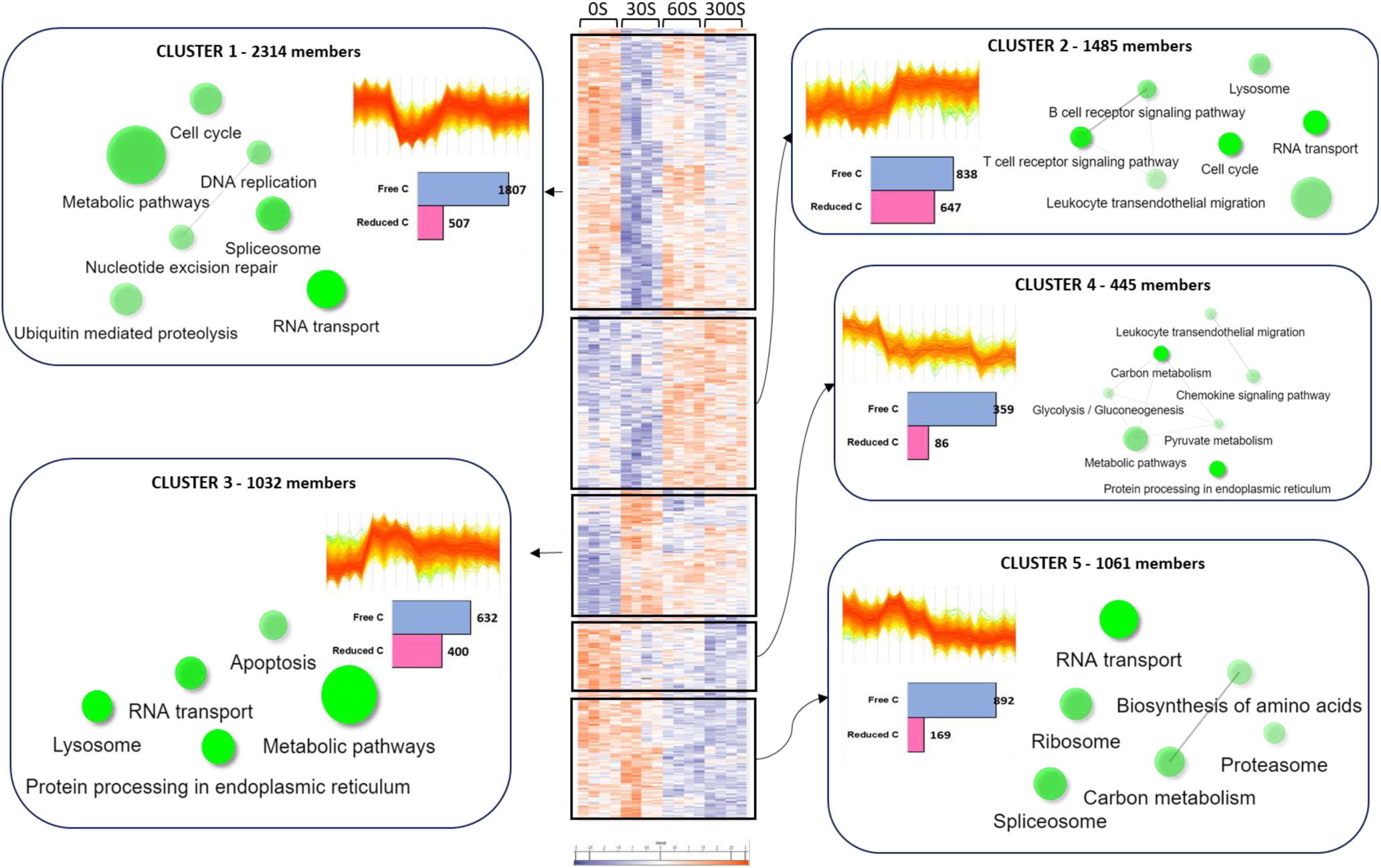
Dynamics of the regulation of freeCys and RmCys containing peptides in T cells after activation from 0 sec to 300 sec in the four conditions. Hierarchical clustering of all regulated Free/RmCys peptides (5813 [free Cys] and 1,560 [RmCys]) reveal a significant modulation of cysteines during T cell activation as early as 30 sec of antigen activation. The regulated peptides could be subdivided into 5 distinct clusters based on their common regulation trends. The information of each sub clusters are further elaborated as five distinct cluster profiles; number of peptides from each fraction is depicted by bar plots. A protein level, sub-cluster wise KEGG pathway (-log(2) FDR) enrichment analysis was performed, resulting in the illustrated pathways extracted from the 5 clusters. The size of the spheres indicate the number of proteins involved in the given term and the colour intensity indicate the relative FDR score. The hierarchical clustering was performed using Z-score clustering in the program Perseus. The KEGG enrichment was performed using the ShinyGo tool.

Cluster 1 was one of the largest and most dynamic clusters, comprising 1,807 freeCys peptides and 507 RmCys peptides, all of which were downregulated within 30 seconds of activation. Downregulation of freeCys indicates that, upon stimulation, these cysteines either become reversibly modified, participate in disulfide bond formation, or conjugate to metal ions. In contrast, downregulation of RmCys peptides reflects an increase in free cysteine states at these sites following stimulation. Proteins represented in Cluster 1 were predominantly involved in metabolic pathways, RNA splicing, and RNA transport, biological processes in which regulatory cysteines have previously been shown to play important roles (37–39).

Consistent with the notion that cysteines can function as molecular switches with bidirectional regulatory potential (40), sub-cluster 3, in contrast to sub-cluster 1, consisted of peptides that were upregulated within 30 seconds yet participated in similar cellular functions, including metabolism and RNA transport. This highlights the dynamic and context-dependent nature of cysteine regulation.

Cluster 2 also contained a substantial number of regulated cysteine-containing peptides (838 freeCys and 647 RmCys) and was notably enriched in proteins associated with the T-cell receptor signaling pathway. In contrast, Cluster 4 comprised proteins for which predominantly freeCys peptides were progressively downregulated across all three stimulation time-points relative to unstimulated cells.

Collectively, the clustering analysis reveals extensive and rapid modulation of cysteine residues across proteins involved in diverse cellular pathways, including DNA/RNA-related processes, metabolism, and T-cell signaling itself. These findings demonstrate that cysteine regulation is highly dynamic even during very brief TCR activation and establish cysteine chemistry as a major regulatory layer in cellular signaling processes underlying T-cell activation (**Figure 3**).

### Transient exposure of disulfide-bonded cysteines reflects conformational and mechanical remodeling at the immunological synapse

Cluster 3 in **Figure 3** include freeCys that show increased accessibility already 30 seconds after activation were strongly enriched in extracellular or luminal domains of disulfide-rich proteins, including integrin subunits, focal adhesion and adhesion regulators, tetraspanin-associated proteins, and secretory-pathway glycoproteins (**Figure 4**). At the site level, regulated cysteines mapped predominantly to flexible loops, domain boundaries, and linker regions within PSI, hybrid, and EGF-like domains-regions known to undergo large-scale conformational rearrangements during receptor activation and mechanical force transmission.

**Figure 4:**
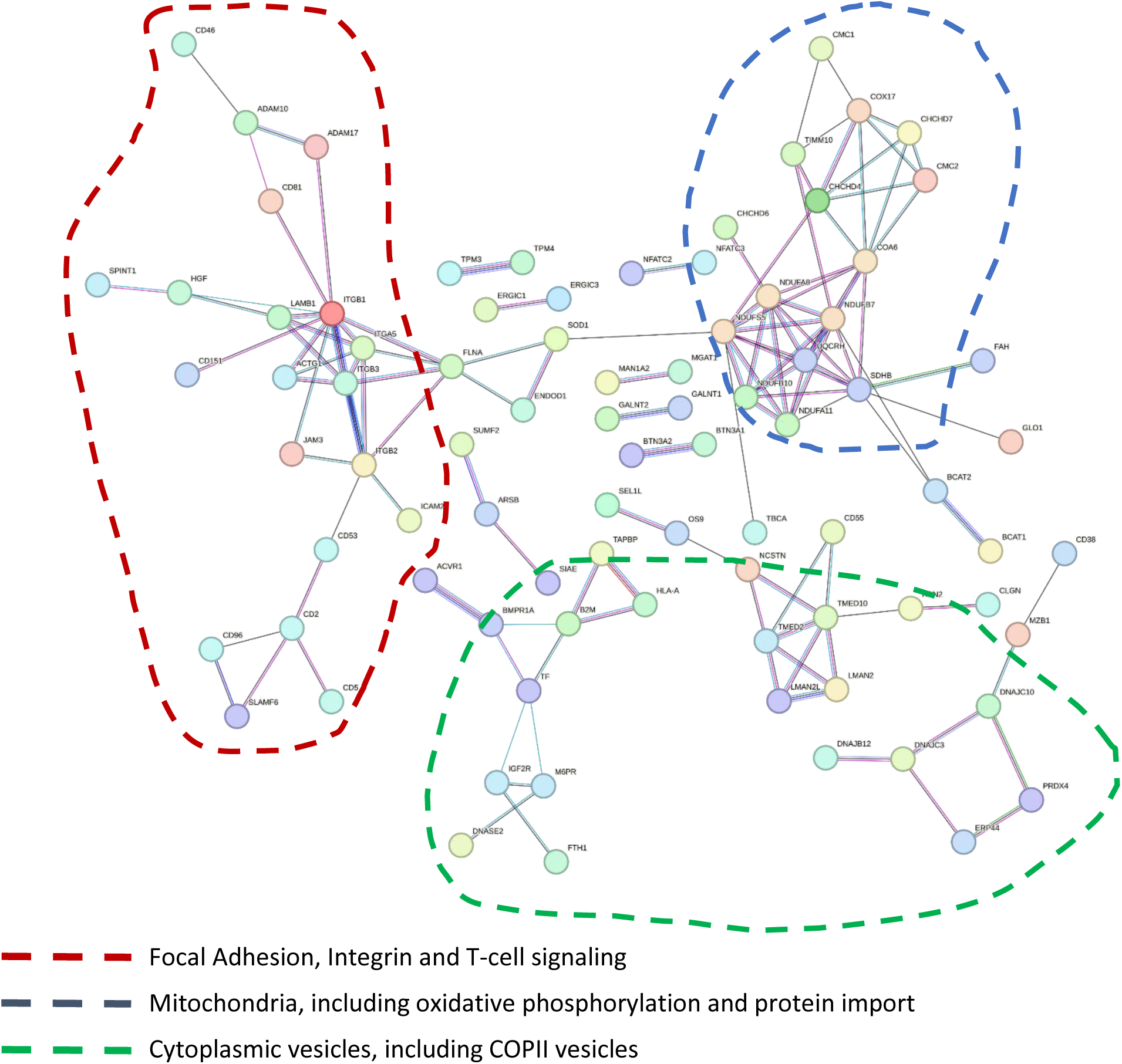
FreeCys upregulated after 30 seconds. A total of 160 proteins contained freeCys that were significantly upregulated after already 30 sec of TCR activation. The lines define proteins involved in the mentioned processes and compartments mentioned in the figure.

The kinetics of these changes revealed both transient and sustained classes of thiol exposure. Many integrin-associated cysteines displayed a rapid increase at 30 seconds followed by partial normalization at later time points, consistent with short-lived conformational states associated with inside-out activation, extension, and clustering. Other cysteines, particularly within luminal secretory-pathway proteins, exhibited more sustained accessibility, suggesting establishment of a new redox or structural steady state during synapse maturation.

Collectively, these observations support a model in which early TCR signaling drives rapid mechanical and organizational changes at the cell surface that transiently expose normally buried cysteines without requiring stable disulfide bond reduction or covalent rearrangement.

### Rapid and selective unexpected regulation of mitochondrial freeCys during early T-cell activation

A striking feature of the early cysteine remodeling landscape was the strong enrichment of mitochondrial proteins among sites showing increased freeCys accessibility already 30 seconds after TCR activation (**Figure 4**). This rapid modulation indicates that mitochondria participate directly in the initiation phase of T-cell signaling, rather than acting solely as downstream metabolic effectors.

Several mitochondrial proteins involved in redox control and metabolic regulation, including superoxide dismutase 1 (SOD1) and enzymes of central metabolic pathways, exhibited pronounced and often transient increases in freeCys accessibility at this early time point. In the case of SOD1, multiple cysteine residues showed concurrent regulation in both free and reversibly modified pools, consistent with rapid redox cycling and dynamic control of antioxidant activity during early signaling.

The predominance of increased freeCys in mitochondrial proteins argues against mitochondrial oxidative stress, which would be expected to reduce free thiols and promote irreversible oxidation. Instead, these changes are consistent with regulated cysteine exposure or reduction, likely mediated by mitochondrial thioredoxin and peroxiredoxin systems that rapidly buffer activation-induced redox fluctuations. In parallel, early TCR signaling is known to induce mitochondrial repositioning toward the immunological synapse, where mitochondria support localized ATP production, calcium buffering, and redox regulation (41). Such spatial reorganization may expose mitochondrial proteins to distinct ionic and redox microenvironments, leading to rapid changes in cysteine accessibility through conformational remodeling or altered protein-protein interactions.

Beyond redox control, several regulated mitochondrial FreeCys were located in enzymes associated with energy metabolism and biosynthetic precursor generation, suggesting a role in early metabolic priming. Cysteine-based regulation of these enzymes provides a fast and reversible mechanism to tune mitochondrial function before transcriptional or translational reprogramming occurs.

Collectively, these observations support a model in which mitochondrial cysteine chemistry acts as an early regulatory layer that integrates redox signaling, metabolic readiness, and spatial organization during the initiation of T-cell activation, operating in parallel with phosphorylation-based signaling cascades.

### Loss of free cysteines in zinc-binding proteins reveals metal coordination as an early regulatory mechanism

In contrast to the widespread increases in cysteine accessibility observed after TCR activation, a distinct subset of freeCys-containing peptides exhibited a pronounced loss of free thiol signal within 30 seconds of stimulation. Specifically, 151 proteins showed a reduction of more than 40% in free cysteine abundance at this early time point. Notably, these downregulated freeCys sites were highly enriched in proteins annotated or predicted to bind zinc, with more than 50% containing cysteine-rich motifs characteristic of zinc finger, RING, LIM, or other metal-coordinating domains.

Zinc is an essential trace element involved in numerous cellular processes and has been shown to play a critical role in T-cell activation (5, 42, 43). Zinc-binding proteins often contain short, cysteine- and histidine-rich motifs that coordinate one or more zinc ions, thereby stabilizing defined protein folds and enabling interactions with DNA, RNA, proteins, or lipids (44). The most prevalent zinc-finger motif in humans is the C2H2-type, in which two cysteines and two histidines are separated by 17–19 amino acids (45).

Across the combined freeCys and RmCys datasets, we quantified 5,981 cysteine-containing peptides corresponding to 951 annotated zinc finger proteins. Among these, 1,140 unique peptides were significantly regulated at one or more time points following TCR activation. Of these regulated peptides, 365 contained CxxC or CxC motifs, consistent with metal coordination. Notably, 253 peptides were detected in the freeCys fraction, whereas 111 peptides were identified in the RmCys fraction. Functional annotation of proteins harbouring these zinc-binding peptides revealed enrichment in gene expression regulation and RNA metabolic processes (**Figure 5A**), molecular functions such as metal binding, DNA/RNA binding, and transferase activity (**Figure 5B**), and predominant localization to the nucleus and intracellular organelles (**Figure 5C**).

**Figure 5:**
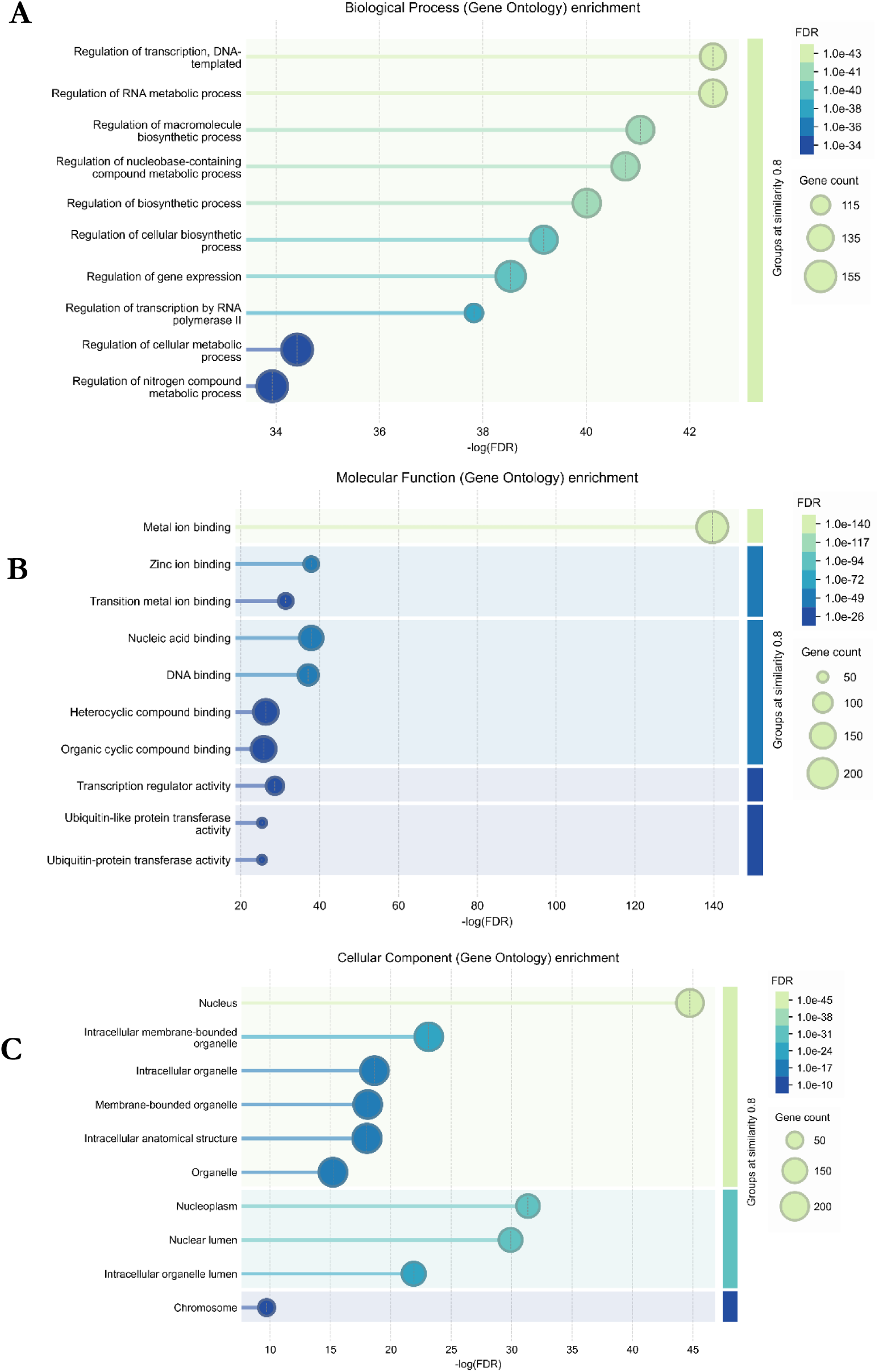
Regulated Zinc finger proteins. A total of 365 of the regulated zinc finger proteins that contained CxxC or CxC motifs were analysed for its (A) Biological function, (B) Molecular functions and (C) Cellular compartments using STRING. The size of the sphere indicate the number of proteins involved and the colour intensity indicate the relative FDR score (p-value cut-off 0.05)

The rapid loss of free cysteine signal at these sites within 30 seconds of T-cell activation is most parsimoniously explained by zinc coordination or stabilization of zinc-binding structures, rather than by oxidative modification. Zinc binding effectively masks cysteine thiols from CysPAT labeling and is well known to induce conformational changes that regulate protein–protein interactions, enzymatic activity, and subcellular localization. The speed and selectivity of these changes suggest that zinc availability and coordination are dynamically regulated during early TCR signaling, potentially through localized zinc fluxes or redistribution triggered by receptor engagement.

Together, these findings identify zinc coordination as a rapid and parallel regulatory mechanism operating alongside phosphorylation and redox-dependent conformational changes, enabling fast functional modulation of signaling, scaffolding, and transcriptional regulatory proteins during early T-cell activation, independent of new protein synthesis.

### Examples of cysteine-dependent regulation of proximal TCR complexes and membrane-associated regulators

Although the CD3 chains themselves are not cysteine-rich in their cytoplasmic ITAM regions, multiple membrane-proximal regulators of TCR signaling exhibit early cysteine modulation, indicating that cysteine chemistry acts immediately upstream and alongside phosphorylation.

LCK, the Src-family kinase responsible for initiating TCR signaling, shows regulation of Cys217 in the freeCys fraction. This residue lies outside the catalytic site but within a region implicated in conformational control and protein–protein interactions. Modulation of free cysteine accessibility at this position within 30 seconds of activation suggests that LCK activity is not only controlled by phosphorylation (e.g., Y394/Y505) but may also be tuned by cysteine-dependent conformational or redox mechanisms during signal initiation.

In parallel, the receptor-type tyrosine phosphatase CD45 (PTPRC) displays dynamic regulation at multiple cysteine sites. Notably, Cys311 and Cys434 in the extracellular domain show opposite temporal behaviour compared with Cys754 and Cys1169 in the cytoplasmic phosphatase-containing region. This domain-specific pattern strongly suggests distinct regulatory mechanisms acting on CD45 across the membrane, potentially coupling extracellular conformational changes to intracellular phosphatase activity. Given CD45’s central role in controlling LCK phosphorylation state, these cysteine dynamics provide a plausible mechanism for rapid, reversible tuning of phosphatase function during early TCR activation.

Downstream of LCK, ZAP-70 is the next essential kinase recruited to phosphorylated ITAMs. In your dataset, ZAP-70 exhibits regulated freeCys changes at Cys78 and Cys84, both located within the SH2 domain responsible for binding phosphorylated CD3 chains. These sites are distinct from the well-characterized Cys39 (not detected due to peptide size constraints) but lie within the same functional module.

Regulation of cysteine accessibility within the SH2 domain is highly significant: it implies that cysteine chemistry may influence SH2-mediated binding and complex stability, potentially affecting the dwell time or orientation of ZAP-70 at the TCR complex. Importantly, this cysteine modulation occurs on the same timescale as robust ZAP-70 phosphorylation in your phosphoproteomics data, supporting a model in which phosphorylation and cysteine-based regulation jointly control ZAP-70 activation and signaling output.

While a key component of the T cell signalosome LAT itself is not strongly cysteine-regulated in the dataset, several LAT-associated adaptors and downstream effectors show clear cysteine modulation. These include proteins involved in assembling and stabilizing the signalosome, where protein–protein interactions rather than catalytic activity dominate.

Cysteine regulation in this layer is characterized mainly by freeCys changes, consistent with conformational rearrangements or altered interaction interfaces rather than stable covalent modification. This supports the idea that cysteine chemistry contributes to dynamic signalosome assembly and disassembly, complementing phosphorylation-dependent recruitment.

PLCγ1, a key effector linking TCR signaling to Ca²⁺ flux and DAG production, exhibits regulated cysteine sites that show delayed but sustained changes, particularly in the freeCys fraction. This temporal pattern differs from the rapid changes observed in LCK and ZAP-70 and suggests a two-phase regulatory mechanism: phosphorylation-driven activation followed by cysteine-based stabilization or tuning of PLCγ1 function during sustained signaling.

Downstream transcriptional regulators influenced by Ca²⁺ and DAG signaling also show cysteine modulation. For example, NFAT family members (e.g., NFATC3) display regulated freeCys sites, consistent with known redox sensitivity of NFAT proteins. These changes likely contribute to controlling transcription factor activation, nuclear translocation, or DNA binding during later phases of TCR signaling.

The nucleotide exchange factor RASGRP1, which links DAG signaling to Ras activation, shows regulated freeCys changes at Cys166. This site-level regulation suggests that cysteine chemistry may influence GEF activity or membrane association, providing an additional regulatory layer controlling Ras–MAPK pathway activation downstream of the TCR.

Similarly, AKT exhibits modulation at Cys297, a residue previously implicated in redox-sensitive regulation of kinase activity. This supports a broader theme in which cysteine-based regulation fine-tunes kinase signaling beyond phosphorylation alone.

## Conclusion

This study reveals a previously unrecognized regulatory layer in early T-cell activation, demonstrating that cysteine chemistry is dynamically and selectively remodeled across the TCR signaling pathway within seconds of receptor engagement. By integrating time-resolved analyses of free and reversibly modified cysteines with quantitative phosphoproteomics, we show that cysteine regulation is not a downstream consequence of signaling but an immediate and coordinated component of signal initiation that operates in parallel with phosphorylation.

Cysteine modulation was detected throughout the TCR signaling cascade, encompassing membrane-proximal kinases and phosphatases (LCK, CD45, ZAP-70), signalosome components, second-messenger generators, and downstream transcriptional regulators. These changes were site-specific, bidirectional, and temporally structured, arguing against nonspecific oxidative stress and instead supporting a model of regulated cysteine exposure, masking, and reversible modification that fine-tunes signaling output. In this framework, phosphorylation provides rapid signal amplification, while cysteine-based mechanisms introduce analog control over protein conformation, interaction stability, and enzymatic activity.

A central and unexpected finding of this study is the identification of zinc coordination as a rapid and widespread regulatory mechanism during early T-cell activation. The selective loss of free cysteines in zinc-binding proteins within 30-60 seconds of stimulation indicates dynamic zinc engagement at cysteine-rich motifs, stabilizing functional protein folds and regulatory complexes without requiring new protein synthesis. This zinc-dependent modulation operates alongside redox-driven cysteine remodeling, together forming a chemically distinct but integrated regulatory axis that shapes early signaling decisions.

Equally striking is the rapid regulation of mitochondrial cysteines, positioning mitochondria as active participants in the initiation of T-cell signaling rather than passive metabolic responders. The predominance of increased free cysteines in mitochondrial proteins suggests regulated redox buffering, conformational remodeling, and metabolic priming occur in parallel with proximal TCR signaling. These findings highlight cysteine chemistry as a key mechanism linking phosphorylation-driven signaling to redox control, metal coordination, and organelle function.

Collectively, this work establishes cysteine-based regulation as a fundamental, fast-acting, and multidimensional signaling language in T-cell activation, complementing and extending the canonical phosphorylation-centric paradigm. By revealing how redox chemistry, zinc coordination, and conformational dynamics are integrated into the earliest moments of immune signaling, this study provides a new conceptual framework for understanding T-cell activation and opens unexplored avenues for therapeutic modulation of immune responses through targeted manipulation of cysteine chemistry.

## Acknowledgement

Funded by Lundbeck foundation (RIMMI project). We thank Prof. Blagoy Blagoev for generously providing the Jurkat E6-1 clone, Zap70 and pY493 Zap70 antibodies. Vladimir Gorshkov for helping with technicalities of mass spec as well as data analysis. This research was financially supported by the Novo Nordic Foundation (No. 95-310-73303-01100), the Lundbeck foundation (R336-2020-1113) and the Villum Center for Bioanalytical Sciences at SDU.

---

**Figure S1:**
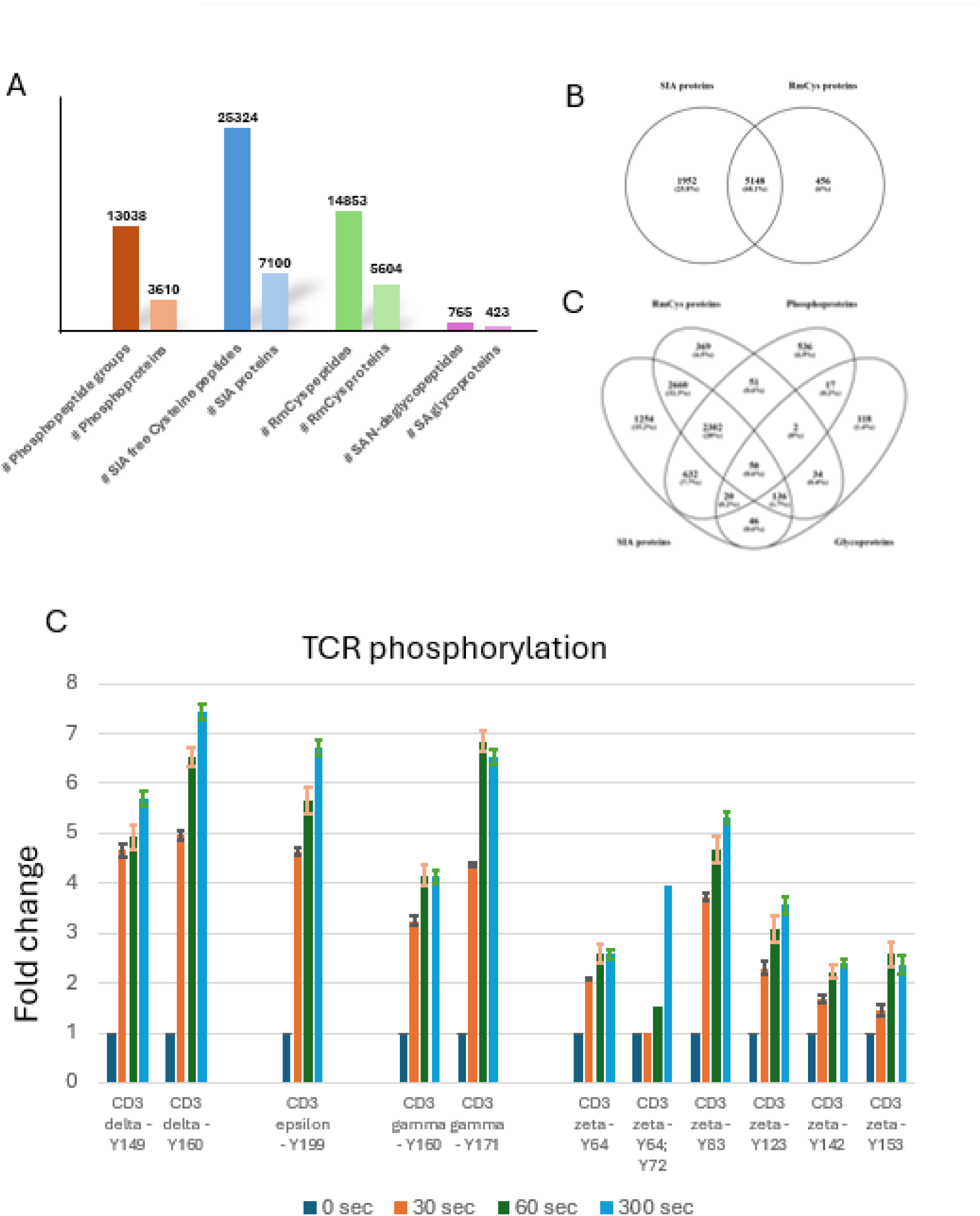
Overview of analysis (A) Identified proteins and unique peptide groups with FreeCys, RmCys, sialic acid containing N-linked glycosylation and phosphorylation, (B) Overlap between RmCys and FreeCys proteins. (C) Overlap between all PTM proteins in the analysis. (D) Regulation of tyrosine-phospho-sites in the CD3 subunits of TCR.

**Figure S2.**
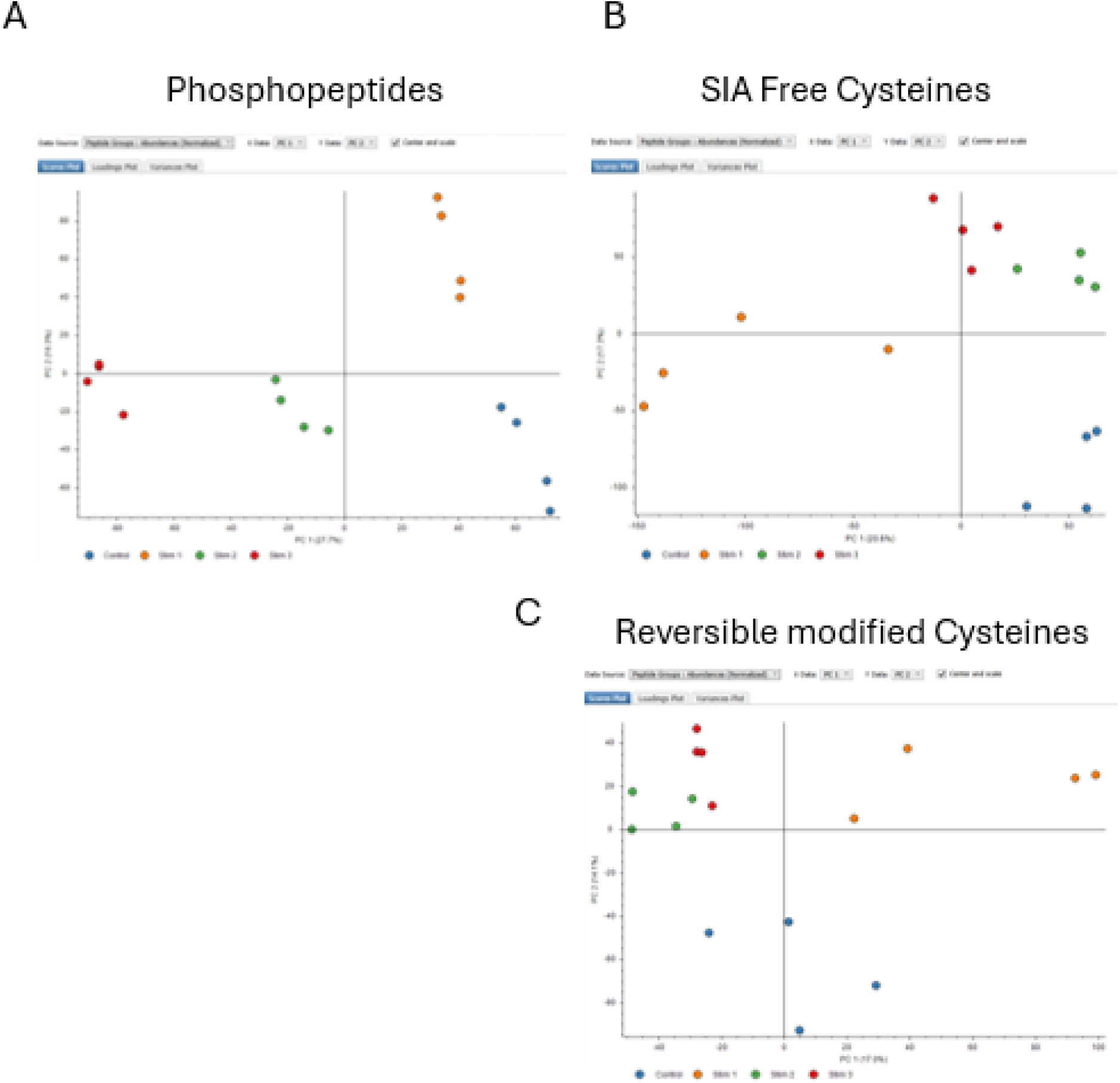
Principal component analysis of (A) Phosphopeptide groups, (B) FreeCys and (C) RmCys

